# Accurate sequence variant genotyping in cattle using variation-aware genome graphs

**DOI:** 10.1101/460345

**Authors:** Danang Crysnanto, Christine Wurmser, Hubert Pausch

**Affiliations:** Animal Genomics, ETH Zurich, Zurich, Switzerland; Chair of Animal Breeding, TU München, Freising, Germany

**Keywords:** Sequence variant genotyping, Genome graph, Variation-aware graph, cattle, Whole-genome sequencing

## Abstract

**Background:** The genotyping of sequence variants typically involves as a first step the alignment of sequencing reads to a linear reference genome. Because a linear reference genome represents only a small fraction of sequence variation within a species, reference allele bias may occur at highly polymorphic or diverged regions of the genome. Graph-based methods facilitate to compare sequencing reads to a variation-aware genome graph that incorporates a collection of non-redundant DNA sequences that segregate within a species. We compared accuracy and sensitivity of graph-based sequence variant genotyping using the *Graphtyper* software to two widely used methods, i.e., *GATK* and *SAMtools*, that rely on linear reference genomes using whole-genomes sequencing data of 49 Original Braunvieh cattle.

**Results:** We discovered 21,140,196, 20,262,913 and 20,668,459 polymorphic sites using *GATK*, *Graphtyper*, and *SAMtools*, respectively. Comparisons between sequence variant and microarray-derived genotypes showed that *Graphtyper* outperformed both *GATK* and *SAMtools* in terms of genotype concordance, non-reference sensitivity, and non-reference discrepancy. The sequence variant genotypes that were obtained using *Graphtyper* had the lowest number of mendelian inconsistencies for both SNPs and indels in nine sire-son pairs with sequence data. Genotype phasing and imputation using the *Beagle* software improved the quality of the sequence variant genotypes for all tools evaluated particularly for animals that have been sequenced at low coverage. Following imputation, the concordance between sequence- and microarray-derived genotypes was almost identical for the three methods evaluated, i.e., 99.32, 99.46, and 99.24% for *GATK*, *Graphtyper*, and *SAMtools*, respectively. Variant filtration based on commonly used criteria improved the genotype concordance slightly but it also decreased sensitivity. *Graphtyper* required considerably more computing resources than *SAMtools* but it required less than *GATK*.

**Conclusions:** Sequence variant genotyping using *Graphtyper* is accurate, sensitive and computationally feasible in cattle. Graph-based methods enable sequence variant genotyping from variation-aware reference genomes that may incorporate cohort-specific sequence variants which is not possible with the current implementations of state-of-the-art methods that rely on linear reference genomes.

## Introduction

The sequencing of important ancestors of many cattle breeds revealed millions of sequence variants that are polymorphic in dairy and beef populations [1–4]. In order to compile an exhaustive catalog of polymorphic sites that segregate in *Bos taurus*, the 1000 Bull Genomes consortium was established [5, 6]. The 1000 Bull Genomes Project imputation reference panel facilitates to infer sequence variant genotypes for large cohorts of genotyped animals thus enabling genomic investigations at nucleotide resolution [5, 7–9].

Sequence variant discovery and genotyping typically involves two steps that are carried out successively [10–13]: first, raw sequencing data are generated, trimmed and filtered to remove adapter sequences and bases with low sequencing quality, respectively, and aligned towards a linear reference genome using, e.g., *Bowtie* [14] or the Burrows-Wheeler Alignment (*BWA)* software [15]. The aligned reads are subsequently compared to the nucleotide sequence of a reference genome in order to discover and genotype polymorphic sites using, e.g., *SAMtools* [16] or the Genome Analysis Toolkit (*GATK)* [17–19]. Variant discovery may be performed either in single- or multi-sample mode. The accuracy (i.e., ability to correctly genotype sequence variants) and sensitivity (i.e., ability to detect true sequence variants) of sequence variant discovery is higher using multi-sample than single-sample approaches particularly when the sequencing depth is low [20–24]. However, the genotyping of sequence variants from multiple samples simultaneously is a computationally intensive task, particularly when the sequenced cohort is large and diverse and had been sequenced at high coverage [19]. The multi-sample sequence variant genotyping approach that has been implemented in the *SAMtools* software has to be restarted for the entire cohort once new samples are added. *GATK* implements two different approaches to multi-sample variant discovery, i.e., the *UnifiedGenotyper* and *HaplotypeCaller* modules, with the latter relying on intermediate files in gVCF format that include probabilistic data on variant and non-variant sites for each sequenced sample. Applying the *HaplotypeCaller* module allows for separating variant discovery within samples from the estimation of genotype likelihoods across samples. Once new samples are added to an existing cohort, only the latter needs to be performed for the entire cohort, thus enabling computationally efficient parallelization of sequence variant genotyping in a large number of samples.

Sequence variant genotyping approaches that rely on alignments to a linear reference genome have limitations to variant discovery, because a haploid reference sequence does not reflect variation within a species. As a result, read alignments may be erroneous particularly at genomic regions that differ substantially between the sequenced individual and the reference sequence, thus introducing reference allele bias, flawed genotypes, and false-positive variant discovery around indels [25–27]. Aligning reads to population- or breed-specific reference genomes may overcome most of these limitations [28–30]. However, considering multiple (population-specific) linear reference genomes with distinct genomic coordinates complicates the biological interpretation and annotation of sequence variant genotypes across populations [31].

Genome graph-based methods consider non-linear reference sequences for variant discovery [31–35]. A variation-aware genome graph may incorporate distinct (population-specific) reference sequences and known sequence variants. Recently, the *Graphtyper* software has been developed in order to facilitate sequence variant discovery from a genome graph that has been constructed and iteratively augmented using variation of the sequenced cohort [32]. So far, sequence variant genotyping using variation-aware genome graphs has not been evaluated in cattle.

An unbiased evaluation of the accuracy and sensitivity of sequence variant genotyping is possible when high confidence sequence variants and genotypes are accessible that were detected using genotyping technologies and algorithms different from the ones to be evaluated [36]. For species where such a resource is not available, the accuracy of sequence variant genotyping may be evaluated by comparing sequence variant to microarray-derived genotypes (e.g., [2, 24]). Due to the ascertainment bias in SNP chip data, this comparison may overestimate the accuracy of sequence variant discovery particularly at variants that are either rare or located in less-accessible genomic regions [37, 38].

In this study, we compare sequence variant discovery and genotyping from a variation-aware genome graph using *Graphtyper* to two state-of-the-art methods (*GATK*, *SAMtools*) that rely on linear reference genomes in 49 Original Braunvieh cattle. We compare sequence variant to microarray-derived genotypes in order to assess accuracy and sensitivity of sequence variant genotyping for each of the three methods evaluated.

## Results

Following quality control (removal of adapter sequences and low-quality bases), we aligned more than 13 billion paired-end reads (2 × 125 and 2 × 150 basepairs) from 49 Original Braunvieh cattle to the UMD3.1 assembly of the bovine genome. On average, 98.44% (91.06-99.59%) of the reads mapped to the reference genome. 4.26% (2.0-10.91%) of the mapped reads were flagged as duplicates and not considered for further analyses. The average sequencing depth per animal was 12.75 and it ranged from 6.00 to 37.78. The average sequencing depth of 31 samples was above 12-fold. Although the re-alignment of sequencing reads around indels is no longer required when sequence variants are genotyped using the latest version of *GATK* (v 4), it is still recommended to improve the genotyping of indels using *SAMtools*. To ensure a fair comparison of the three tools evaluated, we realigned the reads around indels on all BAM files and used the re-aligned files as a starting point for our comparisons (Figure 1). The sequencing read data of 49 cattle were deposited at the European Nucleotide Archive (ENA) (http://www.ebi.ac.uk/ena) under primary accession PRJEB28191.

**Figure 1.**
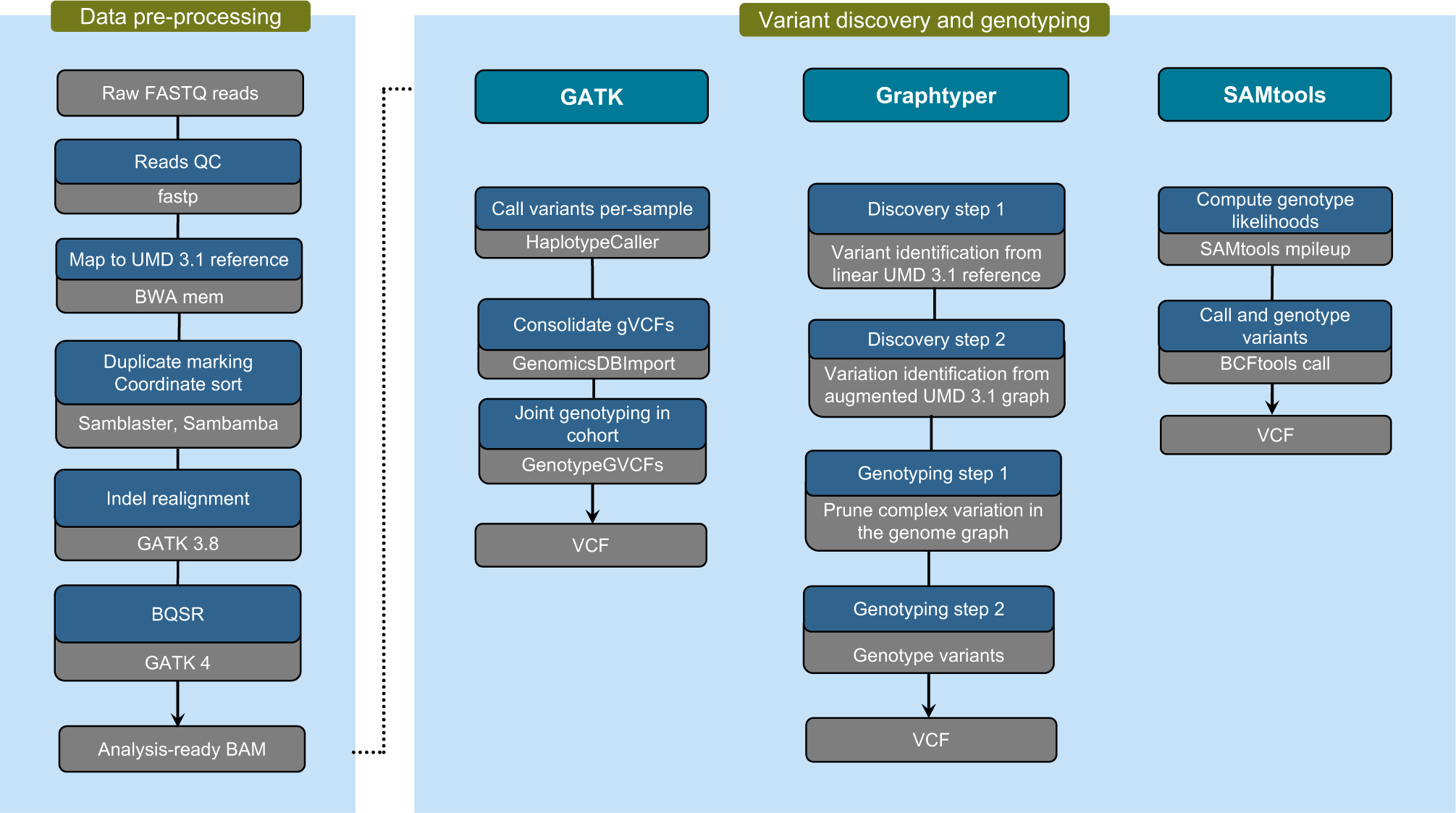
Schematic representation of three sequence variant discovery and genotyping methods evaluated. According to the best practice recommendations for sequence variant discovery using *GATK*, the VQSR module should be applied to distinguish between true and false positive variants. Because this approach requires a truth set of variants which is not (publicly) available for cattle, the VQSR module was not considered in our evaluation.

### Sequence variant discovery and genotyping

Polymorphic sites (SNPs, short insertions and deletions) were discovered and genotyped in the 49 animals using either *GATK* (version 4), *Graphtyper* (version 1.3) or *SAMtools* (version 1.8). All software programs were run using default parameters and workflow descriptions for variant discovery (Figure 1, also see Material and Methods). Only autosomal sequence variants were considered to evaluate the accuracy and sensitivity of sequence variant genotyping. Because variant filtering has a large impact on the accuracy and sensitivity of sequence variant genotyping [39, 40], we evaluated both the raw variants that were detected using default parameters for variant discovery (Figure 1) and variants that remained after applying filtering criteria that are commonly used but may differ slightly between different software tools. Please note that *GATK* was run with filtering parameters suggested when applying *Variant Quality Score Recalibration* (VQSR) is not possible.

Using default parameters for variant discovery for each software program evaluated, the number of polymorphic sites discovered was 21,140,196, 20,262,913, and 20,668,459 using *GATK, Graphtyper* and *SAMtools*, respectively (Table 1). The vast majority (86.79%, 89.42% and 85.11%) of the detected variants were biallelic SNPs. Of the 18,594,182, 18,120,724 and 17,592,038 SNPs detected using *GATK*, *Graphtyper* and *SAMtools*, respectively, 7.46%, 8.31% and 5.02% were novel, i.e., they were not among 102,091,847 polymorphic sites of the most recent version (150) of the Bovine dbSNP database. The transition to transversion (Ti/Tv) ratio of the detected SNPs was 2.09, 2.07 and 2.05 using *GATK*, *Graphtyper* and *SAMtools*, respectively. Using *GATK* revealed four times more multiallelic SNPs (246,220) than either *SAMtools* or *Graphtyper*.

**Table 1.** Numbers of different types of autosomal sequence variants detected in 49 Original Braunvieh cattle using three sequence variant genotyping methods (Full) and subsequent variant filtration based on commonly used criteria (Filtered).

|  | Full |  |  | Filtered |  |  |
| --- | --- | --- | --- | --- | --- | --- |
|  | *GATK* | *Grphtyper* | *SAMtools* | *GATK* | *Grphtyper* | *SAMtools* |
| Variants | 21,140,196 | 20,262,913 | 20,668,459 | 19,761,679 | 17,679,155 | 18,871,549 |
| SNPs | 18,594,182 | 18,120,724 | 17,592,038 | 17,248,593 | 15,777,446 | 16,272,917 |
| Not in dbSNP | 1,387,781 | 1,505,586 | 882,575 | 867,838 | 564,326 | 570,901 |
| Biallelic | 18,347,962 | 18,053,396 | 17,528,249 | 17,111,806 | 15,730,153 | 16,218,714 |
| Multi-allelic | 246,220 | 67,328 | 63,789 | 136,787 | 47,293 | 54,203 |
| Ti/Tv ratio | 2.09 | 2.07 | 2.05 | 2.17 | 2.18 | 2.16 |
| SNP array (%) |  |  |  |  |  |  |
| BovineHD | 99.46 | 99.61 | 99.32 | 99.21 | 98.79 | 98.85 |
| Bovine SNP50 | 99.14 | 99.26 | 99.12 | 98.91 | 98.87 | 98.90 |
| Indels | 2,478,489 | 2,044,585 | 3,076,421 | 2,445,766 | 1,826,808 | 2,598,632 |
| Not in dbSNP | 663,831 | 596,137 | 1,279,162 | 639,219 | 456,752 | 979,291 |
| Biallelic | 2,166,352 | 1,753,391 | 2,704,413 | 2,133,840 | 1,571,195 | 2,310,386 |
| Multi-allelic | 312,137 | 291,194 | 372,008 | 311,926 | 255,613 | 288,246 |
| Insertion/Deletion | 0.88 | 0.88 | 1 | 0.88 | 0.88 | 0.99 |
| Complex variation | 67,525 | 97,604 | 0 | 67,320 | 74,901 | 0 |

The number of indels identified was 2,478,489, 2,044,585, and 3,076,421 using *GATK*, *Graphtyper*, and *SAMtools*, respectively, and 26.78%, 29.15%, and 41.75% of them were novel. *SAMtools* revealed the highest number and proportion (14.9%) of indels. Between 12 and 14% of the detected indels were multiallelic. While *Graphtyper* and *GATK* identified more (12%) deletions than insertions, the ratio was almost equal using *SAMtools*.

On average, each Original Braunvieh cattle carried between 7 and 8 million variants that differed from the UMD3.1 reference genome. Of those, between 2.4 and 2.6 million SNPs were homozygous for the alternate allele, between 3.8 and 4.7 million SNPs were heterozygous and between 0.7 and 1 million were indels (Table 2).

**Table 2.** Average number of autosomal variants identified per animal using three sequence variant genotyping methods. The number of variants is presented for the three tools evaluated before (Full) and after (Filtered) applying recommended filters to identify and exclude low quality variants.

|  | Full |  |  | Filtered |  |  |
| --- | --- | --- | --- | --- | --- | --- |
|  | *GATK* | *Grphtyper* | *SAMtools* | *GATK* | *Grphtyper* | *SAMtools* |
| Total biallelic SNPs | 6,324,455 | 7,384,058 | 6,617,948 | 6,105,674 | 6,533,711 | 6,564,229 |
| Heterozygous | 3,890,351 | 4,758,297 | 4,187,882 | 3,744,336 | 4,074,011 | 4,147,033 |
| Homozygous ALT | 2,434,104 | 2,625,761 | 2,430,066 | 2,361,338 | 2,459,700 | 2,417,196 |
| Ti/Tv | 2.17 | 2.13 | 2.11 | 2.20 | 2.14 | 2.13 |
| Total biallelic indels | 693,697 | 767,261 | 1,007,420 | 691,765 | 697,637 | 960,218 |
| Heterozygous | 390,495 | 441,172 | 616,981 | 388,622 | 391,856 | 593,417 |
| Homozygous ALT | 303,202 | 326,089 | 390,439 | 303,143 | 305,781 | 366,801 |
| Singletons | 49,166 | 23,406 | 32,810 | 41,408 | 17,999 | 32,398 |

An intersection of 15,901,526 biallelic SNPs was common to all sequence-variant discovery tools evaluated (Figure 2A), i.e., between 85.51 and 90.39% of the detected SNPs of each tool, and 466,029 (2.93%, Ti/Tv: 1.81) of them were novel, i.e., they were not present in dbSNP 150. The Ti/Tv-ratio of the common SNPs was 2.22. *SAMtools* had the largest number of SNPs in common with the other two tools (90.39%). The number of private SNPs, i.e., SNPs that were detected by one but not the other tools was highest for *GATK* and lowest for *Graphtyper*.

**Figure 2.**
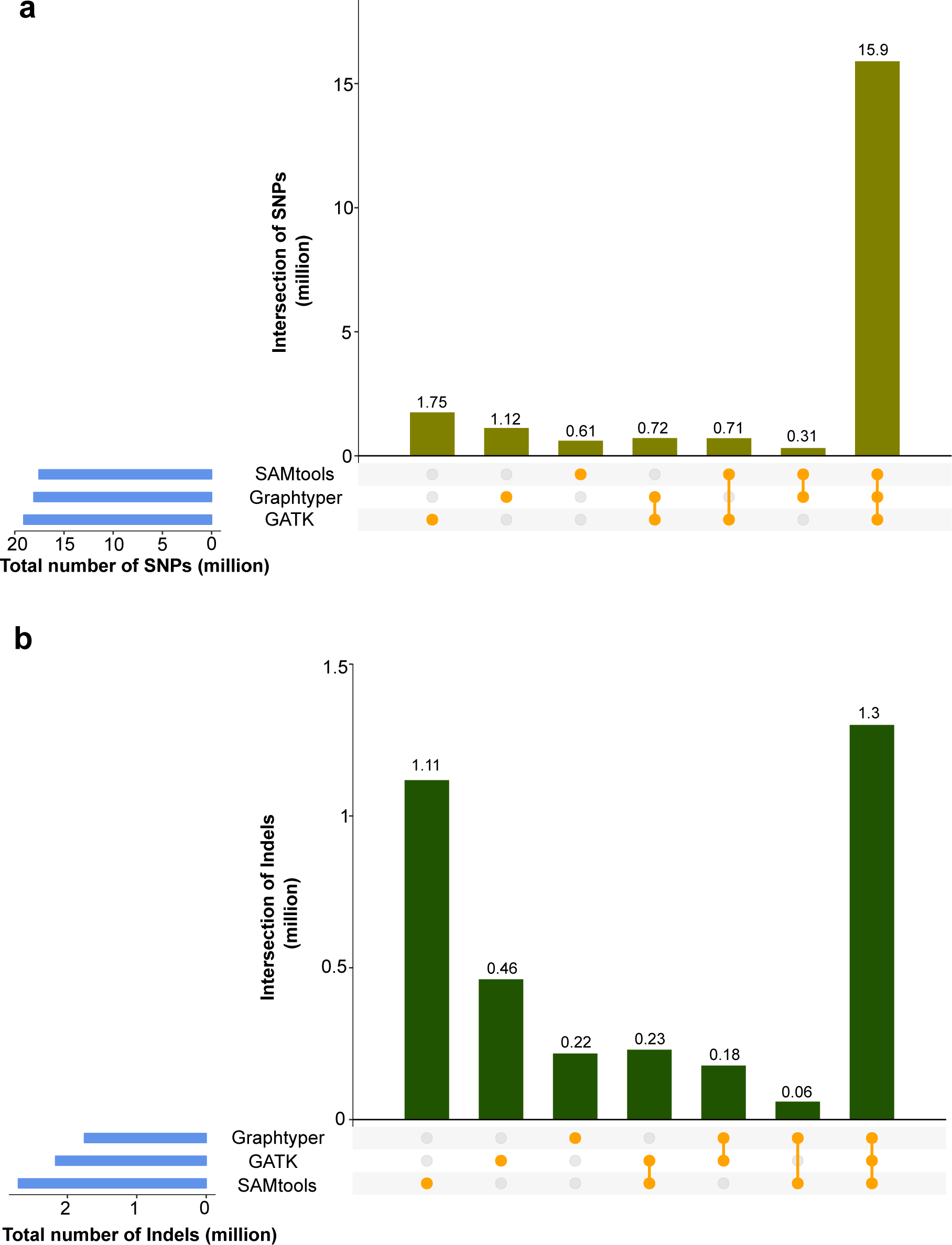
Number of biallelic SNPs (A) and indels (B) identified in 49 Original Braunvieh cattle using three sequence variant genotyping methods. Blue horizontal bars represent the total number of sites discovered for each method. Vertical bars indicate private and common variants detected by the methods evaluated.

The number of biallelic indels (Figure 2B) that was common to all tools evaluated was 1,299,467 and 98,931 (13.13%) of those were novel, i.e., they were not present in dbSNP 150. The intersection of the three tools was considerably smaller for indels than SNPs. *Graphtyper* had the highest proportion of indels in common with the other tools (74.11%). *SAMtools* discovered the highest number (2,704,413) of biallelic indels and most of them (41.26%) were not detected using either *GATK* or *Graphtyper*. *GATK* (21.2%) and *Graphtyper* (12.38%) discovered less private indels than *SAMtools*.

### Sequence variant genotyping using *Graphtyper* is accurate

The 49 sequenced animals were also genotyped using either Illumina BovineHD or Illumina BovineSNP50 Bead chips. Genotype concordance, non-reference sensitivity and non-reference discrepancy were calculated using array-called and sequence variant genotypes at corresponding positions. The genotype concordance is a measure for the proportion of variants that have identical genotypes in the microarray and whole-genome sequencing data. The non-reference sensitivity is the proportion of microarray-derived variants that were also detected in the sequencing data. Non-reference discrepancy reflects the proportion of sequence variants that have genotypes that differ from the microarray-derived genotypes (see Additional file 4 for more details on how the different metrics were calculated). All metrics were calculated both for raw and filtered variants either before or after applying the algorithm implemented in the *Beagle* software for haplotype phasing and imputation.

In the raw data, the proportion of missing non-reference sites was 1.90%, 0.56%, and 0.47% using *GATK*, *Graphtyper*, and *SAMtools*, respectively. The genotype concordance between the sequence- and microarray-derived genotypes was higher (P < 0.005) using *Graphtyper* (97.72%) than either *SAMtools* (97.68%) or *GATK* (95.99%) (Table 3). For the three tools evaluated, the genotype concordance was higher at homozygous than heterozygous sites, particularly in animals that have been sequenced at low depth (Additional file 1). In order to take the variable proportions of missing genotypes in the sequence variants into account, we calculated non-reference sensitivity and non-reference discrepancy. Non-reference sensitivity was almost identical using *Graphtyper* (98.26%) and *SAMtools* (98.21%). However, non-reference sensitivity was clearly lower using *GATK* (93.81%, *P* < 0.001). Non-reference discrepancy was lower using *Graphtyper* (3.53%) than either *SAMtools* (3.6%, *P* = 0.003) or *GATK* (6.35%, *P* < 0.001).

**Table 3.** Comparisons between array-called and sequence variant genotypes. Genotype concordance, non-reference sensitivity and non-reference discrepancy (in percentage) was calculated between the genotypes from the Bovine SNP Bead chip and sequence–derived genotypes for 49 Original Braunvieh cattle considering either the raw or imputed (imp) sequence variant genotypes before (full) and after (filtered) variants were filtered based on commonly used criteria. Asterisks denote that the best value (italic) differs significantly from the other two values (* *P* ≤ 0.05, ** *P* ≤ 0.01, *** *P* ≤ 0.001).

|  | Genotype concordance |  |  |  | Non-reference sensitivity |  |  |  | Non-reference discrepancy |  |  |  |
| --- | --- | --- | --- | --- | --- | --- | --- | --- | --- | --- | --- | --- |
|  | full |  | filtered |  | full |  | filtered |  | full |  | filtered |  |
|  | raw | imp | raw | imp | raw | imp | raw | imp | raw | imp | raw | imp |
| *GATK* | 95.99*** | 99.32*** | 96.02*** | 99.39*** | 93.81*** | 99.36 | 93.67*** | 99.15 | 6.35*** | 1.05*** | 6.3*** | 0.95*** |
| *Graphtyper* | 97.71 | 99.46 | 97.75 | 99.52 | 98.26 | 99.35 | 97.91 | 99.00*** | 3.53 | 0.83 | 3.47 | 0.73 |
| *SAMtools* | 97.68*** | 99.24*** | 97.7* | 99.29*** | 98.21 | 99.35 | 97.53*** | 98.67*** | 3.6** | 1.17*** | 3.56** | 1.09*** |

Next, we analysed the proportion of opposing homozygous genotypes for SNPs and indels in nine sire-son pairs that were included among the sequenced animals (Table 4). We observed that SNPs that were discovered using either *Graphtyper* or *SAMtools* had almost a similar proportion of genotypes with mendelian inconsistencies in the full and filtered datasets, whereas the values were two times higher using *GATK*. The proportion of opposing homozygous genotypes was higher for indels than SNPs for all tools evaluated. However, it was lower using *Graphtyper* than either *GATK* or *SAMtools* in the full and filtered datasets. Using filtering parameters that are commonly applied for the three tools evaluated (see Material & Methods), we excluded 1,378,517 (6.52%, Ti/Tv 1.24), 2,583,758 (12.75%, Ti/Tv 1.47) and 1,796,910 (8.69%, Ti/Tv 1.36) variants due to low mapping or genotyping quality from the *GATK*, *Graphtyper*, and *SAMtools* datasets, respectively. The genotype concordance between sequence- and microarray-derived genotypes was slightly higher in the filtered than raw genotypes, but the non-reference sensitivity was less in the filtered than raw genotypes indicating that the filtering step also removed some true variant sites from the raw data (Table 3). The filtering step had barely an effect on the proportion of mendelian inconsistencies detected in nine sire-son pairs (Table 4).

**Table 4.** Proportions of opposing homozygous genotypes observed in nine sire-son pairs. The ratio (in percentage) was calculated using autosomal sequence variants considering either the raw or imputed (imp) sequence variant genotypes before (full) and after (filtered) variants were filtered based on commonly used criteria. Asterisks indicate that the best value (italic) differs significantly from the other two values (* *P* ≤ 0.05, ** *P* ≤ 0.01, *** *P* ≤ 0.001).

|  | SNPs |  |  |  | indels |  |  |  |
| --- | --- | --- | --- | --- | --- | --- | --- | --- |
|  | full |  | filtered |  | full |  | filtered |  |
|  | raw | imp | raw | imp | raw | imp | raw | imp |
| *Bovine HD SNP array* | 0.001 |  |  |  |  |  |  |  |
| *GATK* | 0.73* | 0.15* | 0.72* | 0.13* | 0.98* | 0.24* | 0.99* | 0.21* |
| *Graphtyper* | 0.36 | 0.11 | 0.36 | 0.11 | 0.54 | 0.13 | 0.54 | 0.13 |
| *SAMtools* | 0.33 | 0.28* | 0.32 | 0.25* | 0.67 | 0.54* | 0.61 | 0.57* |

### Beagle genotype refinement improved the genotype quality

We used the *Beagle* software to refine the primary genotype calls and infer missing genotypes in the raw and filtered datasets. Following imputation, the quality of the sequence variant genotypes increased for all tools evaluated particularly in cattle where the sequencing coverage was less than 12-fold (Figure 3). The largest increase in the concordance metrics was observed for the sequence variants obtained using *GATK* (Table 3 & 4). Following imputation, the variants identified using *Graphtyper* had significantly higher quality (*P < 0.05*) in eight out of ten metrics evaluated.

**Figure 3.**
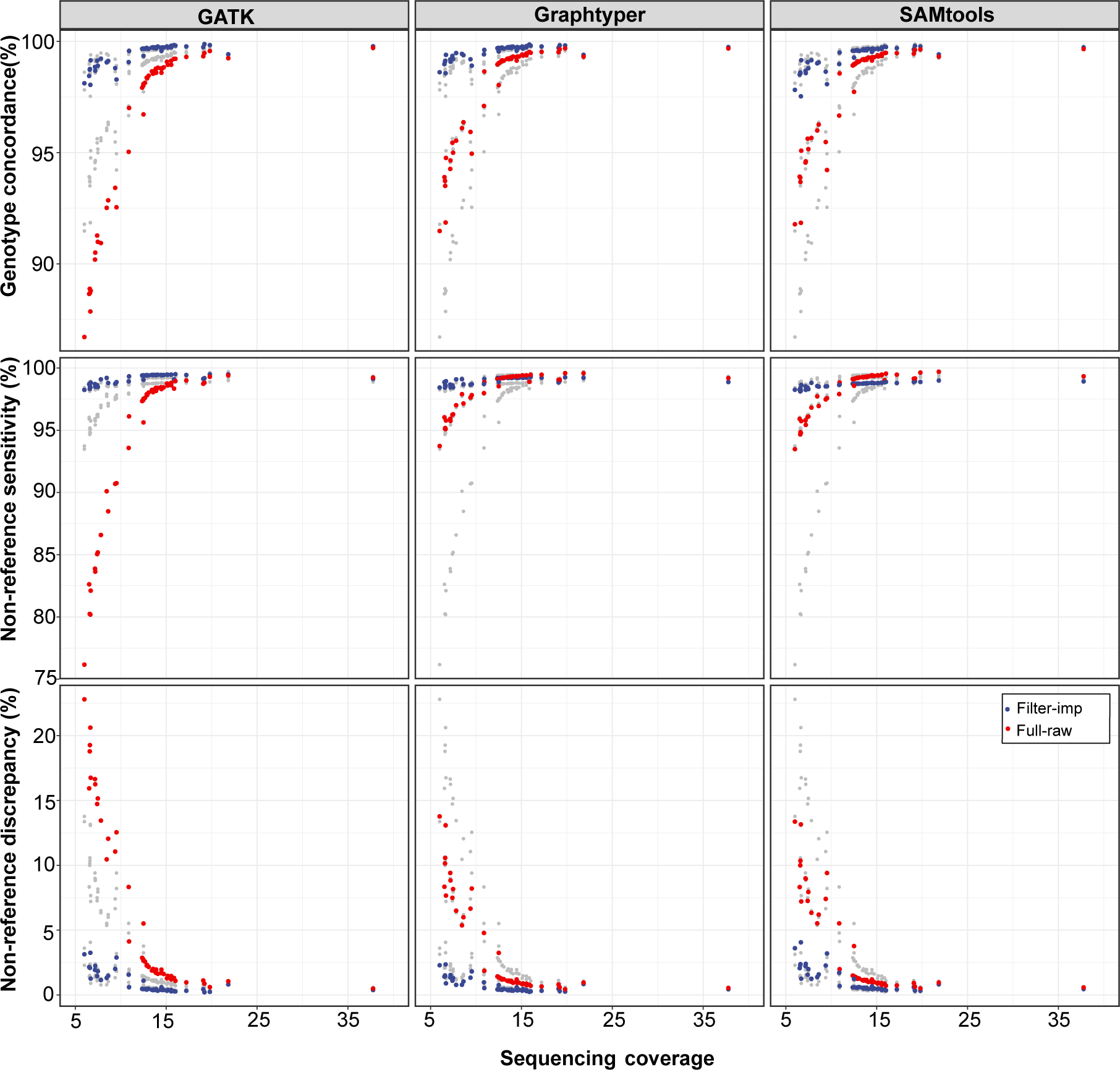
Accuracy and sensitivity of sequence variant genotyping at different sequencing depth. Genotype concordance, non-reference sensitivity and non-reference discrepancy were calculated for 49 Original Braunvieh cattle considering either raw (red) or filtered and imputed (blue) sequence variant genotypes. The grey points represent overlays of the results from the other methods.

The quality of the sequence variant genotypes, particularly before *Beagle* genotype phasing and imputation, was influenced by the various depth of coverage in the 49 sequenced samples of our study (Figure 3). When we restricted the evaluations to 31 samples with an average sequencing depth above 12-fold, the three tools performed almost identical (Additional file 5). However, *Graphtyper* was significantly (*P < 0.05)* better in 12 (out of total 20 metrics) than either *GATK* or *SAMtools*. When 18 samples with an average sequencing depth less than 12-fold were considered, the differences observed in the three metrics were more pronounced between the three tools. In samples with low sequencing coverage, *Graphtyper* performed significantly (*P < 0.05)* better than either *GATK* or *SAMtools* in all concordance metrics both before and after filtering and *Beagle* imputation except for the non-reference sensitivity.

### Computing requirements

The multi-sample sequence variant genotyping pipelines that were implemented using either *GATK* or *SAMtools* were run separately for each chromosome in single-threading mode. The *SAMtools mpileup* module took between 3.07 and 11.4 CPU hours and it required between 0.12 and 0.25 gigabytes (GB) peak random-access memory (RAM) per chromosome. To genotype 20,668,459 sequence variants in 49 animals, *SAMtools* mpileup required 192 CPU hours (Figure 4).

**Figure 4.**
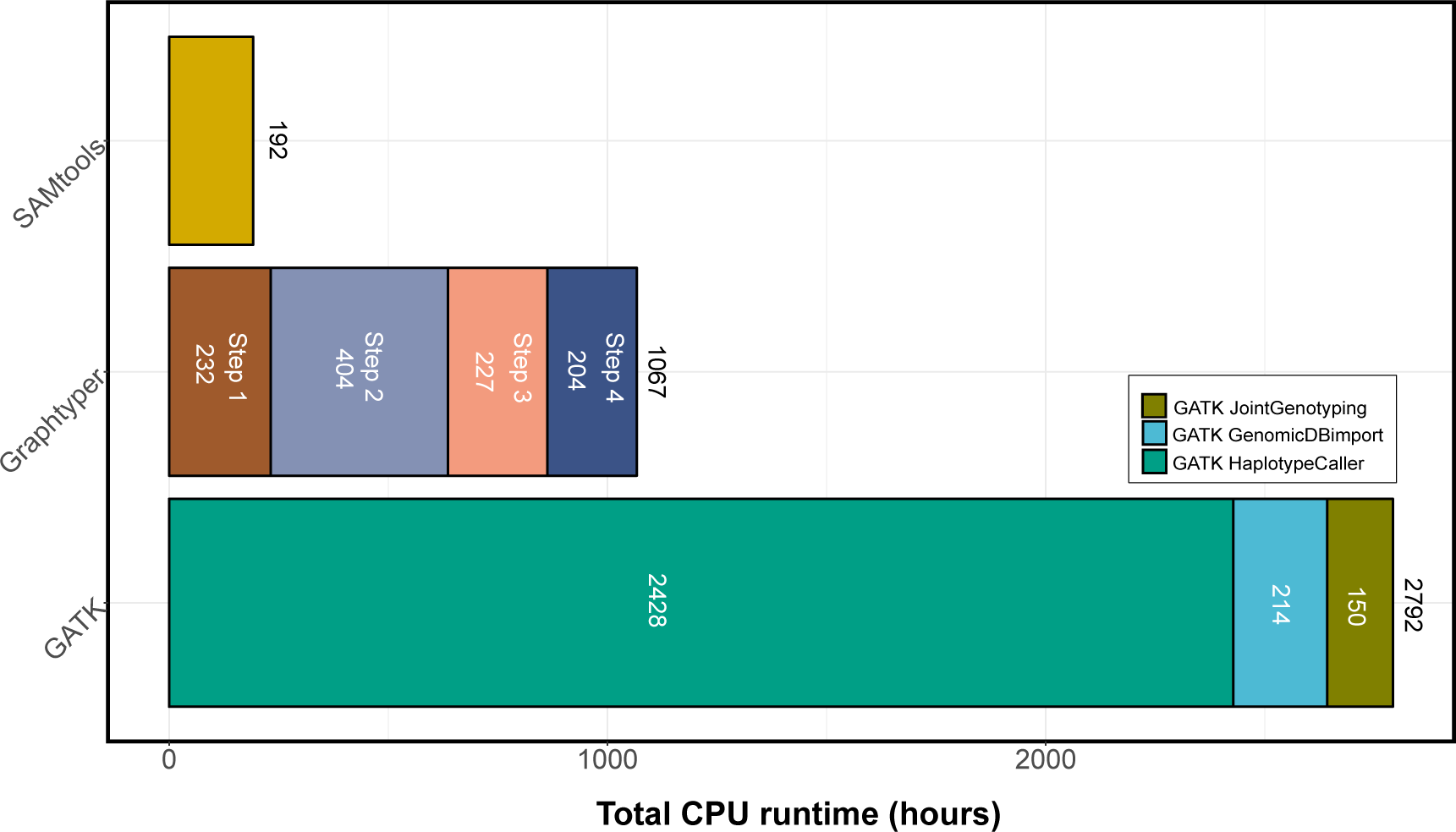
Computing time required to genotype all autosomal sequence variants in 49 Original Braunvieh cattle. The runtime of *GATK* and *Graphtyper* is shown for the different steps (see Figure 1 for more details).

For *GATK*, we submitted 1421 parallel jobs of the *HaplotypeCaller* module (*i.e.*, one job for each animal and chromosome) that required between 3.9 and 12.3 GB RAM and between 0.36 and 11 CPU hours to complete. To process 29 chromosomes in 49 samples, the *HaploytpeCaller* module required a total of 2428 CPU hours. Subsequently, we ran the *GATK GenomicsDBImport* module which required between 7.98 and 20.88 GB RAM and between 2.81 and 19.31 CPU hours per chromosome. *GATK Joint Genotyping* required between 4.33 and 17.32 GB of RAM and between 1.81 and 14.01 hours per chromosome. To genotype 21,140,196 polymorphic sequence variants in 49 animals, the *GATK* pipeline required a total of 2792 CPU hours (Figure 4).

The *Graphtyper* pipeline including the construction of the variation graph and the genotyping of sequence variants was run in parallel for 2538 non-overlapping segments of 1 million base-pairs as recommended by Eggertson *et al.* [32]. The peak RAM required by *Graphtyper* was between 1 and 3 GB per segment. Twelve segments for which *Graphtyper* either ran out of memory or did not finish within the allocated time were subdivided into smaller segments of 10 Kb and subsequently re-run (Additional file 2). The genotyping of 20,262,913 polymorphic sites in 49 animals using our implementation of the *Graphtyper* pipeline took a total of 1066 CPU hours (Figure 4).

The computing resources required by *SAMtools* and *GATK* increased linearly with chromosome length. The computing time required to genotype sequence variants was highly heterogeneous along the genome using *Graphtyper*. The average CPU time per 1 Mb segment was 0.42 hours, but it varied in between 0.196 and 10.11 hours. We suspected that flaws in the reference genome might increase the complexity of the variation-aware graph and that the construction of the graph might benefit from an improved assembly. To test this hypothesis, we re-mapped the sequencing reads to the recently released new bovine reference genome (ARS-UCD1.2, https://www.ncbi.nlm.nih.gov/assembly/GCF_002263795.1) and repeated the graph-based sequence variant discovery. We indeed observed a decrease in the computing time required to genotype polymorphic sites (particularly at chromosomes 12, 27 and 29) and a more uniform runtime along the genome possibly indicating that graph-based variant discovery in cattle will be faster and more accurate from highly contiguous reference sequences (Figure 5).

**Figure 5.**
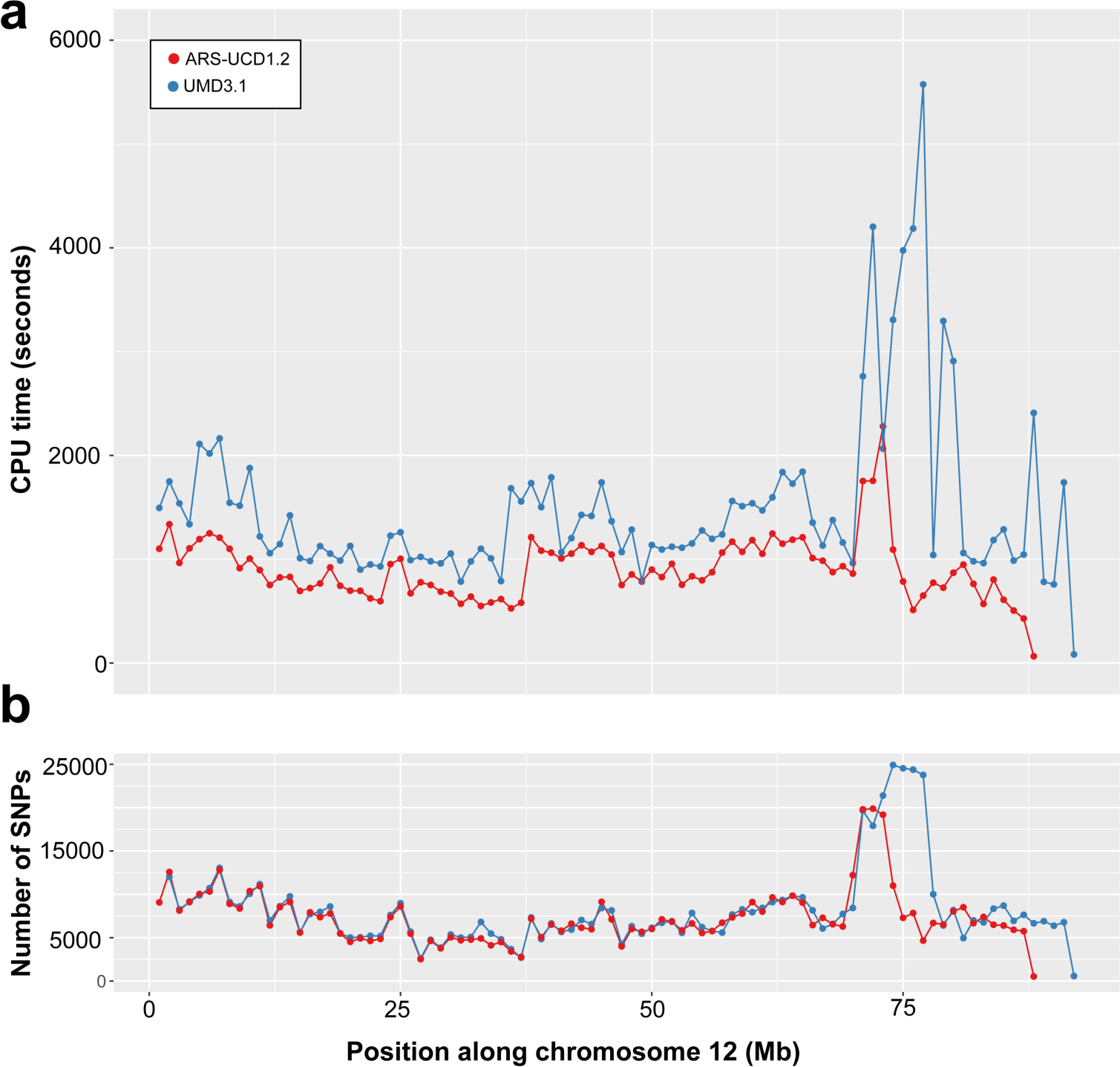
Sequence variant genotyping on chromosome 12 using Graphtyper. Computing time required (A) and number of variants discovered (B) for bovine chromosome 12 using *Graphtyper*. Each dot represents an interval of 1 million basepairs. Blue and red colour represents values for the UMD3.1 and ARS-UCD1.2 versions of the bovine assembly.

## Discussion

We used either *Graphtyper, SAMtools, or GATK* to discover and genotype polymorphic sequence variants in whole genome sequencing data of 49 Original Braunvieh cattle that have been sequenced at between 6 and 38-fold genome coverage. While *SAMtools* and *GATK* discover variants from a linear reference genome, *Graphtyper* locally re-aligns reads to a variation-aware reference graph that incorporates cohort-specific sequence variants [32]. Our graph-based variant discovery pipeline that has been implemented using the *Graphtyper* software used the existing bovine reference sequence to construct the genome graph. The graph was subsequently augmented with variants that were detected from linear alignments of the 49 Original Braunvieh cattle. The use of more sophisticated genome graph-based approaches that have been developed very recently facilitates to map raw sequencing reads directly against a genome graph without the need to first align reads towards a linear reference [34]. While genome graph-based variant discovery has been explored recently in mammalian-sized genomes [27, 31, 32, 35], our work is the first to apply graph-based sequence variant genotyping in cattle.

In order to evaluate graph-based variant discovery in cattle, we compared accuracy and sensitivity of *Graphtyper* to *SAMtools* and *GATK*, i.e., two state-of-the-art methods on linear reference genomes that have been evaluated thoroughly in many species including cattle [2, 20]. We ran each tool with default parameters for variant discovery and applied commonly used or recommended filtration criteria. However, our evaluation of the software tools may suffer from ascertainment bias because we relied on SNPs that have been included in Bovine SNP arrays, i.e., they are located predominantly at well accessible regions of the genome [37, 38, 41]. Thus, the global accuracy and sensitivity of sequence variant discovery might be overestimated in our study. However, this ascertainment bias is unlikely to affect the relative performance of the methods evaluated.

In 49 Original Braunvieh cattle, sequence variant genotyping was more accurate using *Graphtyper* than either *GATK* or *SAMtools.* The difference in accuracy is small between the three tools particularly when samples are sequenced at an average coverage higher than 12-fold (Additional file 5). Yet, *Graphtyper* performed significantly better than *GATK* and *SAMtools* for samples sequenced at medium (>12-fold) or low (<12-fold) coverage indicating that genome graph-based variant discovery in cattle is accurate across a wide range of sequencing depth. *GATK* might perform better than observed in our study when the VQSR module is applied to train the variant filtration algorithm on true and false variants [42]. However, to the best of our knowledge, the required sets of true and false variants are not available in cattle. An intersection of variants detected by different sequence variant genotyping software may be considered as a truth set (e.g., [43]) and compiling such a set is possible using the 49 samples from our study. However a truth set that has been constructed from the data that are used for evaluation is likely to be depleted for variants that are difficult to discover in the target data set, thus preventing an unbiased evaluation of variant calling [36]. Variants from the 1000 Bull Genomes project [5, 6] could potentially serve as a truth/training set. However, variants from the 1000 Bull Genomes project were detected from short read sequencing data using either *GATK* or *SAMtools*, i.e., technologies and software that are part of our evaluation, thus precluding an unbiased comparison of variant discovery between *GATK*, *SAMtools* and *Graphtyper* [36]. Vander Jagt *et al.* [44] showed in a subset of samples from the 1000 Bull Genomes Project that VQSR has barely an effect on the concordance metrics calculated in our study. Interestingly, the proportion of opposing homozygous genotypes in sire/offspring pairs was slightly higher in their study using *GATK* VQSR than *GATK* hard-filtering as used by the 1000 Bull Genomes Project [44]. Considering that the quality of the truth/training sets has a large impact on the capabilities of VQSR and that high-confidence variants are currently not publicly available for cattle, we ran *GATK* using filtering parameters recommended when VQSR is not possible.

Irrespective of the method evaluated, we observed heterozygous under-calling in animals that have been sequenced at low coverage, i.e., heterozygous variants were erroneously genotyped as homozygous due to an insufficient number of sequencing reads supporting the heterozygous genotype [10, 45–47]. In agreement with previous studies [2, 5], *Beagle* imputation improved genotype concordance and reduced heterozygous under-calling particularly in cattle that had been sequenced at low coverage. After the imputation step, the genotype concordance, non-reference sensitivity, and non-reference discrepancy of the three tools was almost identical, indicating that genotyping sequence variants from samples with medium coverage is possible at high accuracy (at least for common variants in more-accessible regions of the genome) using either of the three tools evaluated and subsequent *Beagle* error correction. While such conclusions have been drawn previously for *SAMtools* and *GATK* [2, 20], our findings demonstrate that the genotype likelihoods estimated from the *Graphtyper* software are also compatible with and benefit from the imputation algorithm implemented in the *Beagle* software. Considering that sequence data are enriched for rare variants that are harder to impute than common variants from SNP microarrays [48], the benefits from *Beagle* error correction might be overestimated in our study. An integration of phasing and imputation of missing genotypes directly in a graph-based variant genotyping approach would simplify sequence variant genotyping from variation-aware graphs [31, 49, 50].

Using *Graphtyper* for variant genotyping and *Beagle* for genotype refinement enabled us to genotype sequence variants in 49 Original Braunvieh cattle at a genotypic concordance of 99.52%, i.e., higher than previously achieved using either *GATK* or *SAMtools* for variant calling in cattle that had been sequenced at similar genome coverage [2–5, 20, 51], indicating that graph-based variant discovery might improve sequence variant genotyping. However, applying the filtration criteria that have been recommended for *Graphtyper* [32] removed more variants from the *Graphtyper* (12.75%) than either *GATK* (6.52%) or *SAMtools* (8.69%) datasets. Fine-tuning of the variant filtering parameters is necessary to further increase the accuracy and sensitivity of sequencing variant genotyping, particularly for *Graphtyper* [39, 40]. Moreover, the accuracy and sensitivity of graph-based variant discovery may be higher when known variants are considered for the initial construction of the variation graph [32]. Indeed, we observed a slight increase in genotype concordance (Additional file 3) when we used *Graphtyper* to genotype sequence variants from a variation-aware genome-graph that incorporated bovine variants listed in dbSNP 150. However, additional research is required to prioritize a set of variants to augment bovine genome graphs for different cattle breeds [52].

Using microarray-derived genotypes as a truth set may overestimate the accuracy of sequence variant discovery particularly at variants that are rare or located in less accessible regions of the genome. Moreover, it does not allow for assessing accuracy and sensitivity of indel discovery because variants other than SNPs are currently not routinely genotyped with commercially available microarrays. Estimating the proportion of opposing homozygous genotypes between parent-offspring pairs may be a useful diagnostic metric to detect sequencing artefacts or flawed genotypes at indels [53]. Our results show that genotypes at indels are more accurate using *Graphtyper* than either *SAMtools* or *GATK* because *Graphtyper* produced less opposing homozygous genotypes at indels in nine sire-son pairs than the other methods both in the raw and filtered datasets. These findings are in line with Eggertsson *et al.* [32], who showed that the mapping of the sequencing reads to a variation-aware graph could improve read alignment nearby indels, enabling highly accurate sequence variant genotyping also for variants other than SNPs. Recently, Garrison *et al.* [34] showed that graph-based variant discovery may also mitigate reference allele bias. An assessment of reference allele bias was, however, not possible in our study because the sequencing depth was too low for most samples.

In our study, *Graphtyper* required less computing time than *GATK* to genotype sequence variants for 49 cattle. *SAMtools* required the least computing resources likely because the implemented *mpileup* algorithm produces genotypes from the aligned reads without performing the computationally intensive local re-alignment of the reads. However, with an increasing number of samples, the multi-sample variant genotyping implementation of the *GATK HaplotypeCaller* module seems to be more efficient than *SAMtools mpileup* because variant discovery within samples can be separated from the joint genotyping across samples [19, 44]. A highly parallelized graph-based variant discovery pipeline also offers a computationally feasible and scalable framework for variant discovery in thousands of samples [32]. However, the computing time for graph-based variant genotyping might be high in genomic regions where the nucleotide diversity is high or the assembly is flawed [35, 54]. In our study, the algorithm implemented in the *Graphtyper* software failed to finish within the allocated time for twelve 1 Mb segments including a segment on chromosome 12 that contains a large segmental duplication [48, 55, 56] possibly because many mis-mapped reads increased graph complexity. The region on chromosome 12 contains an unusually large number of sequence variants and has been shown to suffer from low accuracy of imputation [48]. *Graphtyper* also failed to finish within the allocated time for a region on chromosome 23 encompassing the bovine major histocompatibility complex which is known to be rich in diversity. Our results show that *Graphtyper* may also produce genotypes for problematic segments when they are split and processed in smaller bits. Moreover, most of these problems disappeared when we considered the latest assembly of the bovine genome, possibly corroborating that more complete and contiguous genome assemblies may facilitate more reliable genotyping from variation-aware graphs [37, 57].

## Conclusion

Genome graphs facilitate sequence variant discovery from non-linear reference genomes. Sequence variant genotyping from a variation-aware graph is possible in cattle using *Graphtyper*. Sequence variant genotyping at both SNP and indels is more accurate and sensitive using *Graphtyper* than either *SAMtools* or *GATK*. The proportion of mendelian inconsistencies at both SNP and indels is low using *Graphtyper* indicating that sequence variant genotyping from a variation-aware genome graph facilitates accurate variant discovery at different types of genetic variation. Considering highly-informative variation-aware genome graphs that have been constructed from multiple breed-specific de-novo assemblies and high-confidence sequence variants facilitates more accurate, sensitive and unbiased sequence variant genotyping in cattle.

## Methods

### Animal Selection

We selected 49 Original Braunvieh OB bulls that were either frequently used in artificial insemination or explained a large fraction of the genetic diversity of the active breeding population. Semen straws of the bulls were purchased from an artificial insemination center and DNA was prepared following standard DNA extraction protocols.

### Sequencing data pre-processing

All samples were sequenced at either Illumina HiSeq 2500 (30 animals) or Illumina HiSeq 4000 (19 animals) instruments using 150 basepair paired-end sequencing libraries with insert sizes between 400 and 450 basepairs. Quality control (removal of adapter sequences and bases with low quality) of the raw sequencing data was carried out using the *fastp* software (version 0.19.4) with default parameters [58]. The filtered reads were mapped to the UMD3.1 version of the bovine reference genome [59] using *BWA mem* (version 0.7.12) [15] with option-M to mark shorter split hits as secondary alignments, otherwise the default parameters were applied. Optical and PCR duplicates were marked using *Samblaster* (version 0.1.24) [60]. The output of *Samblaster* was converted into BAM format using *SAMtools view* (version 1.3) [16] and subsequently coordinate sorted using *Sambamba* (vesion 0.6.6) [61]. We used the *GATK* (version 3.8) *RealignerTargetCreator* and *IndelRealigner* modules to realign reads around indels. The realigned BAM files served as an input for *GATK* base quality score recalibration using 102,092,638 unique positions from the Illumina BovineHD SNP chip and Bovine dbSNP version 150 as known variants. The *mosdepth* software (version 0.2.2) [62] was used to extract the number of reads covering a genomic position.

### Sequence variant discovery

We followed the best practice guidelines recommended for variant discovery and genotyping using *GATK* (version 4.0.6) with default parameters for all commands [17, 18, 24]. First, genotype likelihoods were calculated separately for each sequenced animal using *GATK HaplotypeCaller* [19], resulting in files in gVCF (genomic Variant Call Format) format for each sample [63]. The gVCF files from 49 samples were consolidated using *GATK GenomicsDBImport*. Subsequently, *GATK GenotypeGVCFs* was applied to genotype polymorphic sequence variants for all samples simultaneously.

*Graphtyper* (version 1.3) was run in multi-sample mode as recommended in Eggertsson *et al.* [32]. Because the original implementation of *Graphtyper* is limited to the analysis of the human chromosome complement, we cloned the *Graphtyper GitHub* repository (https://github.com/DecodeGenetics/graphtyper), modified the source code to allow analyzing the cattle chromosome complement and compiled the program from the modified source code (see Additional file 6). The *Graphtyper* workflow consisted of four steps that were executed successively. First, sequence variants were identified from the read alignments that were produced using *BWA mem* (see above). Next, these cohort-specific variants were used to augment the UMD3.1 reference genome and construct the variation-aware genome graph. Subsequently, the sequencing reads were locally re-aligned against the variation-aware graph. A clean variation graph was produced by removing unobserved haplotypes paths from the raw graph. In the final step, genotypes were identified from the re-aligned reads in the clean graph. The *Graphtyper* pipeline was run in segments of 1 million base-pairs and whenever the program failed to genotype variants for a particular segment either because it run out of memory or exceeded the allocated runtime of twelve hours, the interval was subdivided into smaller segments (10 kb).

Our implementation of *SAMtools mpileup* (version 1.8) [64] was run in multi-sample mode to calculate genotype likelihoods from the aligned reads for all samples simultaneously. The parameters *-E* and *–t* were used to recalculate (and apply) base alignment quality and produce per-sample genotype annotations, respectively. Next, the estimated genotype likelihoods were converted into genotypes using *BCFtools call* using the *-v* and *-m* flags to output variant sites only and permit that sites may have more than two alternative alleles, respectively.

We implemented all pipelines using *Snakemake* (version 5.2.0) [65]. The scripts for the pipelines are available via *GitHub* repository (https://github.com/danangcrysnanto/Graph-genotyping-paper-pipelines).

### Sequence variant filtering and genotype refinement

The *GATK VariantFiltration* module was used to parse and filter the raw VCF files. Quality control on the raw sequencing variants and genotypes was applied according to guidelines that were recommended for each variant caller. Variants that were identified using *GATK* were retained if they met the following criteria: QualByDepth (QD) > 2.0, FisherStrand < 60.0, RMSMappingQuality (MQ) > 40.0, MappingQualityRankSumTest (MQRankSum) > 12.5, ReadPosRankSumTest (ReadPosRankSum) > −8.0, SOR < 3.0 (SNPs) and QD > 2.0, FS < 200.0, ReadPosRankSum > 20.0, SOR < 10.0 (indels). For the variants identified using *SAMtools*, the thresholds that have been applied by the 1000 bull genomes project [5] were considered to remove variants with indication of low quality. Variants were retained if they met the following criteria: QUAL > 20, MQ > 30, ReadDepth (DP) > 10, DP < median(DP) + 3*mean(DP). Moreover, SNPs were removed from the data if they had the same positions as the starting position of an indels. The output of *Graphtyper* was filtered to only include variants that met the criteria recommended by Eggertsson *et al.* [32]: ABHet < 0.0 | ABHet > 0.33, ABHom < 0.0 | ABHom > 0.97, MaxAASR > 0.4, and MQ > 30.

We used *Beagle* (version 4.1) [66] to improve the raw sequence variant genotype quality and impute missing genotypes. The genotype likelihood (*gl*) mode of *Beagle* was applied to infer missing and modify existing genotypes based on the phred-scaled likelihoods (*PL*) of all other non-missing genotypes of the 49 Original Braunvieh animals of our study.

### Evaluation of sequence variant genotyping

To ensure consistent variant representation across the different sequence variant genotyping methods evaluated, we applied the *vt normalize* software (version 0.5) [67]. Normalized variants are parsimonious (i.e., represented by as few nucleotides as possible) and left aligned [67]. The number of variants detected and Ti/Tv ratios were calculated using *vt peek* [67] and *BCFtools stats* [64]. The intersection of variants that were common to the tools evaluated was calculated and visualized using *BCFtools isec* [64] and the UpSet R package [68], respectively.

Mendelian inconsistencies were calculated as the proportion of variants showing opposing homozygous genotypes in nine parent-offspring pairs that were included among the 49 sequenced animals. For this comparison, we considered only sites where the genotypes of both sire and son were not missing.

All 49 sequenced cattle were also genotyped using either Illumina BovineHD (N = 29) or BovineSNP50 (N = 20) Bead chips comprising 777,962 and 54,001 SNPs, respectively. The average genotyping rate at autosomal SNPs was 98.91%. In order to assess the quality of sequence variant genotyping, the genotypes detected by the different variant calling methods were compared to the array-called genotypes in terms of genotype concordance, non-reference sensitivity and non-reference discrepancy [24, 41], see Additional file 4 for more details on the metrics. Non-parametric Kruskal-Wallis tests followed by pairwise Wilcoxon signed-rank tests were applied to determine if either of the three metrics differed significantly between the three tools evaluated.

### Computing environment and statistical analysis

All computations were performed on the ETH Zurich Leonhard Open Cluster with access to multiple nodes equipped with 18 cores Intel Xeon E5-2697v4 processors (base frequency rated at 2.3 GHz) and 128 GB of random-access memory. Unless otherwise stated, the R (version 3.3.3) software environment [69] was used for statistical analyses and ggplot2 (version 3.0.0) [70] was used for data visualisation.

## Declarations

### Animal ethics statement

Not applicable

### Author contributions

Analyzed data: DC HP; Wrote the software pipelines: DC; Generated sequencing data: CW; Wrote the paper: DC HP; Read and approved the final version of the paper: DC CW HP

## Acknowledgements

We thank Braunvieh Schweiz for providing pedigree and genotype data of Original Braunvieh cattle. Semen samples of the sequenced bulls were obtained from Swissgenetics. Sequencing of the Original Braunvieh bulls was partially funded from the Federal Office for Agriculture (FOAG), Bern, Switzerland. The sequence data were partly generated at the Functional Genomics Center Zurich.

## Availability of supporting data

The scripts for three pipelines are available via *GitHub* repository (https://github.com/danangcrysnanto/Graph-genotyping-paper-pipelines). Instructions to install a modified *Graphtyper* version for the bovine chromosome complement are provided in Additional file 6. Sequencing read data of 49 Original Braunvieh bulls are available at the European Nucleotide Archive (ENA) (http://www.ebi.ac.uk/ena) under primary accession PRJEB28191.

## Additional files

### Additional file 1

File format: pdf

Description: Concordance of heterozygous and alternate homozygous genotypes. (a) The average concordance across 49 Original Braunvieh cattle and the concordance at the different sequencing depth based on the (b) raw and (c) imputed datasets.

### Additional file 2

File format: xlsx

Description: Twelve 1 Mb regions where *Graphtyper* initially failed to genotype sequence variants because the algorithm either ran out of memory or exceeded the allocated runtime (12 hours). *Graphtyper* eventually produced genotypes for the sequence variants when these regions were re-run in 10 kb segments.

### Additional file 3

File format: tiff

Description: Accuracy and sensitivity of sequence variant genotyping on bovine chromosome 25 from a variation-aware genome graph that incorporated 2,143,417 dbSNP variants as prior known variants.

### Additional file 4

File format: tiff

Description: Properties of the different metrics used for the evaluation of sequence variant genotyping accuracy. The metrics were calculated using the sum of the red cells as numerator and the cells within the green frame as denominator.

### Additional file 5

File format: xlsx

Description: Sequence variant genotyping quality for 18 and 31 animals that had been sequenced at less and more than 12-fold sequencing coverage, respectively. Asterisks indicate that the best value (italic) differs significantly from the other two values (* *P* ≤ 0.05, ** *P* ≤ 0.01, *** *P* ≤ 0.001).

### Additional file 6

File format: pdf

Description: Description to compile a *Graphtyper* version modified for the cattle chromosome complement.

