## Additional file 1 for "Accurate sequence variant genotyping in cattle using variation-aware genome graphs"

**a**

| | Heterozygous concordance | | | | Homozygous concordance | | | |
|---|---|---|---|---|---|---|---|---|
| | full | | filtered | | full | | filtered | |
| | raw | imp | raw | imp | raw | imp | raw | imp |
| *GATK* | 89.17 | 99.11 | 89.24 | 99.21 | *98.74* | 99.18 | *98.75* | 99.27 |
| *Graphtyper* | *95.79* | *99.36* | *95.82* | *99.44* | 98.55 | *99.51* | 98.59 | *99.57* |
| *SAMtools* | 95.73 | 98.91 | 95.77 | 98.99 | 98.46 | 99.37 | 98.48 | 99.41 |

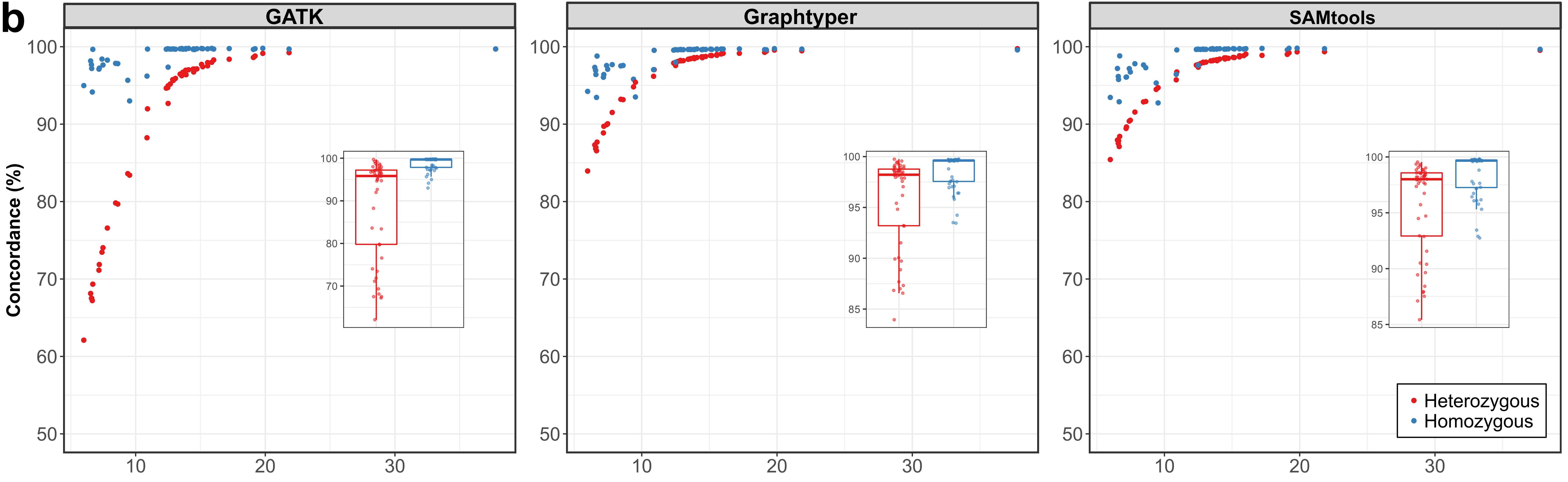

b
GATK
Graphtyper
SAMtools
Concordance (%)
100
90
80
70
60
50
10
20
30
Heterozygous
Homozygous

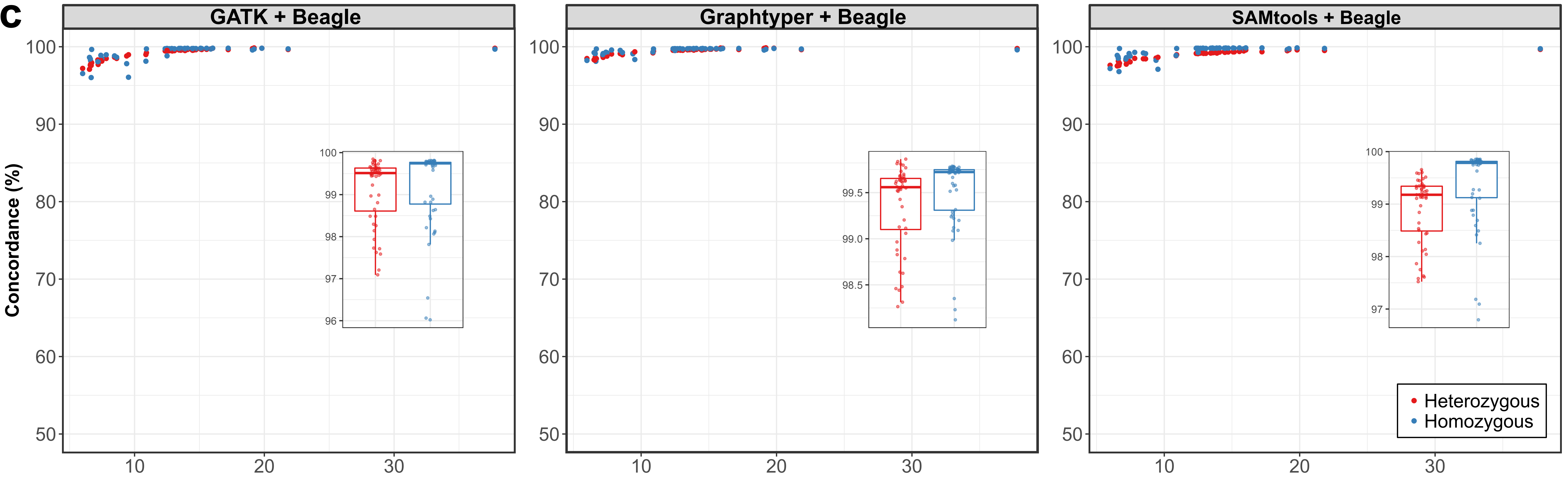

C
GATK + Beagle
Graphtyper + Beagle
SAMtools + Beagle
Concordance (%)
Heterozygous
Homozygous
