## Supplementary figures and images for "Accurate sequence variant genotyping in cattle using variation-aware genome graphs"

### Additional file 3

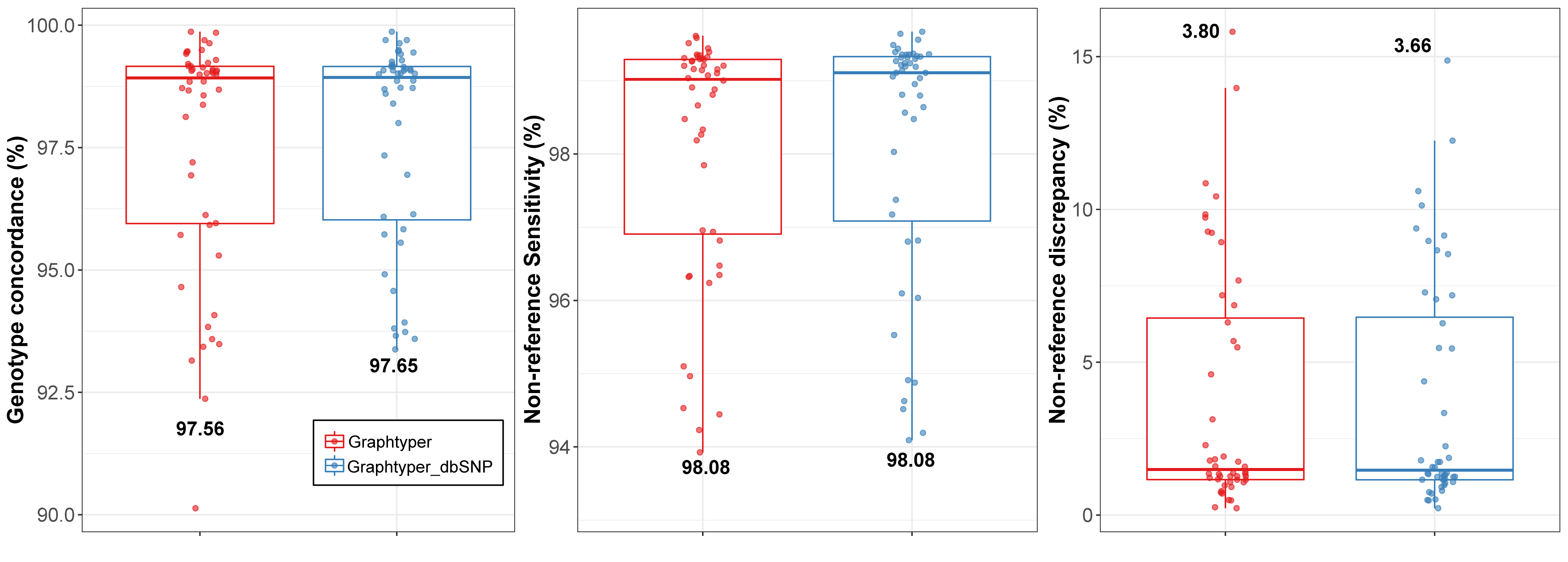

### Additional file 4

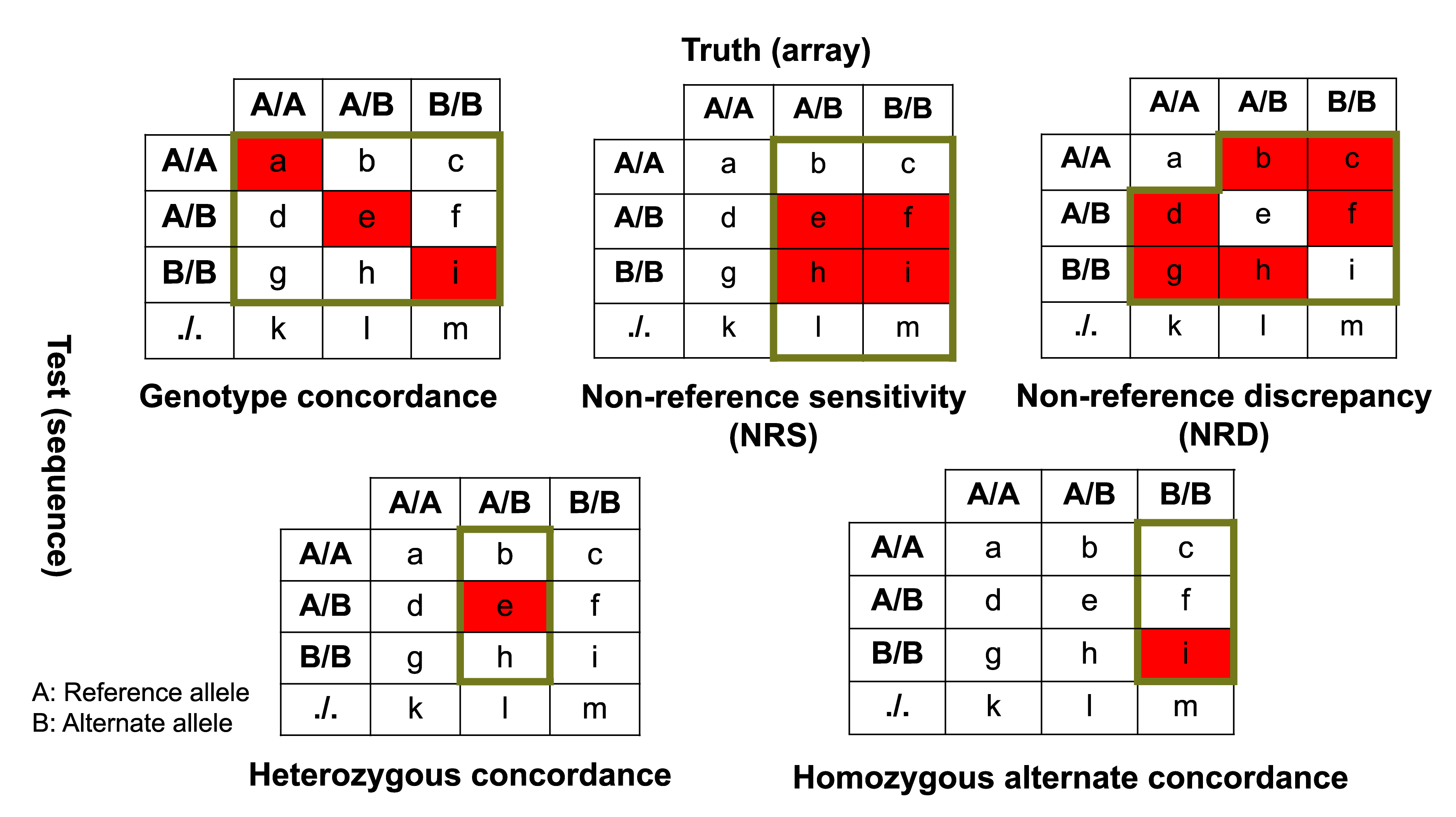
