## Additional file 6 for "Accurate sequence variant genotyping in cattle using variation-aware genome graphs"

### Modified *Graphtyper* for variant discovery and genotyping in cattle

The most convenient way to run a *Graphtyper* version compiled for the bovine chromosome complement is to use Docker (which deals with all required dependencies). The command below starts to download modified *Graphtyper* software hosted at the Dockerhub:

```
docker run --rm cdanang/graphtyper_cattle graphtyper
```

We built the docker images using *Ubuntu* 18.04 as a base image. If you are working on a Linux 64-bit machine you could also get a static executable with command below. We placed the *Graphtyper* binary in /usr/local/bin) and executing command below will copy the *Graphtyper* binary from docker images to the current working directory:

```
docker run --rm -v ${PWD}:/io cdanang/graphtyper_cattle \
cp /usr/local/bin/graphtyper /io

### And then run the software as a standard binary
./graphtyper
```

If you prefer to modify and build a modified version of Graphtyper for the bovine chromosome complement directly from the source, please follow the instructions below:

1. Clone the *Graphtyper Github* at this [link]

   ```
   git clone --recursive https://github.com/DecodeGenetics/graphtyper.git
   ```

2. Create a new `branch` at this specific commit tag. We built graphtyper at this specific commit hash (04ab5ee460fa36129fb0d8ea5d4b72adc3836f52), to compile at the same software version that we use in the paper, please use this commit tag. We named the `branch` as *cattle modification*

   ```
   git checkout -b cattle_modification \
   04ab5ee460fa36129fb0d8ea5d4b72adc3836f52
   ```

3. Change directory into *graphtyper/* and modify the chromosomal specifications in the files *include/graphtyper/graph/absolute_position.hpp* & *src/typer/vcf.cpp* using UMD 3.1 cattle chromosomal names and lengths. The first modification enables all cattle chromosomes (esp. for chromosome number > 23) as the current software release set the maximum allowed length for each chromosomes according to the human *GRChb37 and GRCh38*. The second modifications are required that the respective chromosomal information is written to the *vcf header*.

4. Make sure that these dependencies are installed:

- C++ compiler with C++11 supported (we tested gcc 4.8.5 or gcc 6.3.0 to build the software)
- Boost>=1.57.0
- zlib>=1.2.8
- libbz2
- liblzma
- Autotools, Automake, libtool, Make, and CMake>=2.8.8

5. Follow installation procedures as below. This will put the software in *release-build/bin/graphtyper*

```
mkdir -p release-build && cd release-build
cmake ..
make -j4 graphtyper
bin/graphtyper # Run Graphtyper with modified cattle chromosome
specifications
```
